# Detection of Spatially Aberrant Cells in Spatial Transcriptomics Data by Conformal Prediction

**DOI:** 10.64898/2026.08.12.744448

**Authors:** Zheng Zhang, Xubin Zheng, Qiuyue Yuan, Rui Luo

**Author notes:** Contributing authors.

## Abstract

The hexagonal organization of epithelial cells represents a fundamental feature of normal tissue architecture, reflecting the precise spatial coordination that underlies healthy biological structure. Disruptions to this organization—manifesting as spatially aberrant spots with abnormal gene expression and misplaced positioning—are closely associated with disease initiation and progression. Here, we introduce SPADE, a computational framework that integrates single-cell RNA sequencing and spatial transcriptomics data to quantitatively characterize and detect spatial aberrancy. SPADE leverages a variational autoencoder coupled with Gaussian mixture modeling for cell-type embedding and spatial deconvolution, and incorporates conformal prediction to enable uncertainty-calibrated identification of aberrant spots. Through extensive validation, SPADE demonstrates superior performance in identifying biologically meaningful aberrant spots.

## Introduction

aberrant cell typically refer to cells that significantly deviate from normal state, and are commonly observed in various pathological conditions such as cancer, inflammation, autoimmune diseases, and neurodegenerative disorders [1, 2, 3, 4]. In healthy tissues, morphogen gradients—spatial concentration profiles of signaling molecules—play a central role in guiding cells toward specific fates according to their positional context [5]. Perturbations in these gradients, accompanied by abnormal gene expression programs, can disrupt normal tissue organization and contribute to disease onset and progression. Accordingly, in this study, we characterize aberrant cells by their concurrent deviation from expected spatial positioning and transcriptional dysregulation.

Recent advances in single-cell RNA sequencing (scRNA-seq) have enabled transcriptomic profiling at unprecedented resolution; however, scRNA-seq lacks spatial context, limiting our understanding of how these cellular states integrate within intact tissue architecture. Spatial transcriptomics (ST) technologies extend transcriptomic profiling into the spatial dimension [6, 7, 8, 9], allowing researchers to examine how gene expression and cellular identities are organized across tissues. Despite this progress, most next-generation sequencing (NGS)-based ST methods operate at multi-cellular resolution, where each spot captures transcripts from a mixture of neighboring cells rather than individual cells [10]. This limitation poses challenges in accurately mapping single-cell states to a specific location and detecting subtle spatial deviations. Integration of sc-RNA-seq with ST data presents a promising solution to these challenges.

The rapid accumulation of ST data has motivated the development of diverse computational methods for data integration and analysis. These methods can be broadly classified into two categories based on their primary goals. The first category aims to generate robust spatial embeddings within individual tissue sections to enhance downstream analyses such as spatial domain identification, with graph based methods including SpaMask [11] and GraphST [12] autoencoder based approaches like SEDR [13] and AESTETIK [14], and NMF based methods such as EDGES [14] and STORM [15]. The second category focuses on multi slice integration and batch correction across sections, individuals, or platforms, with SpaBatch [11] using triplet learning for cross slice alignment, SpaCross [16] incorporating a cross masked graph autoencoder with adaptive spatial semantic graph structures, and STAligner [17] offering a GNN based solution for batch corrected embedding generation and consensus domain identification across datasets.

Currently, only a few computational methods have been developed to identify spatially aberrant spots. For instance, the STANDS framework integrates scRNA-seq, ST [18], and histological images to identify aberrant regions by modeling normal spatial expression patterns and detecting regions with high reconstruction error as spatially aberrant domains. In such score-based methods, reconstruction errors serve as anomaly scores, and defining appropriate thresholds is critical for balancing sensitivity and specificity. However, these approaches typically lack a principled mechanism for threshold calibration, which can lead to inconsistent detection performance across datasets.

Here, we propose an uncertainty-calibrated prediction approach (SPADE) to identify aberrant spots by explicitly quantifying uncertainty in spatial positioning. SPADE maps scRNA-seq cells and ST spots into a shared latent space using a variational autoencoder (VAE)-based framework [19]. Within this framework, cellular embeddings are modeled using a Gaussian mixture model (GMM) [20] to capture underlying cell-type structure. Representation of each spatial spot is decomposed into a weighted combination of GMM component centers via non-negative sparse regression, enabling accurate spot deconvolution and mitigating batch effects between scRNA-seq and ST data. SPADE then learns a mapping from spot embeddings to spatial coordinates to estimate the expected spatial position of each spot. Finally, by incorporating a conformal prediction strategy [21], SPADE produces statistically calibrated prediction intervals for each spot’s location, allowing robust and interpretable identification of spatially aberrant cells with controlled false discovery rates.

## Result

### Overview of SPADE

SPADE is a computational framework for identifying spatially aberrant spots by jointly leveraging scRNA-seq and spatial transcriptomics (ST) data. The core principle of SPADE is to learn the mapping rule from transcriptomic to spatial location in normal tissue organization, and to flag spots that deviate from this learned spatial rule. SPADE operates in four stages as shown in Figure 1: (i) cluster-aware latent representation learning, (ii) alignment of cells and spots into a shared latent space, (iii) spatial mapping of ST profiles, and (iv) uncertainty-calibrated aberrance scoring (Fig. 1).

**Fig. 1.**
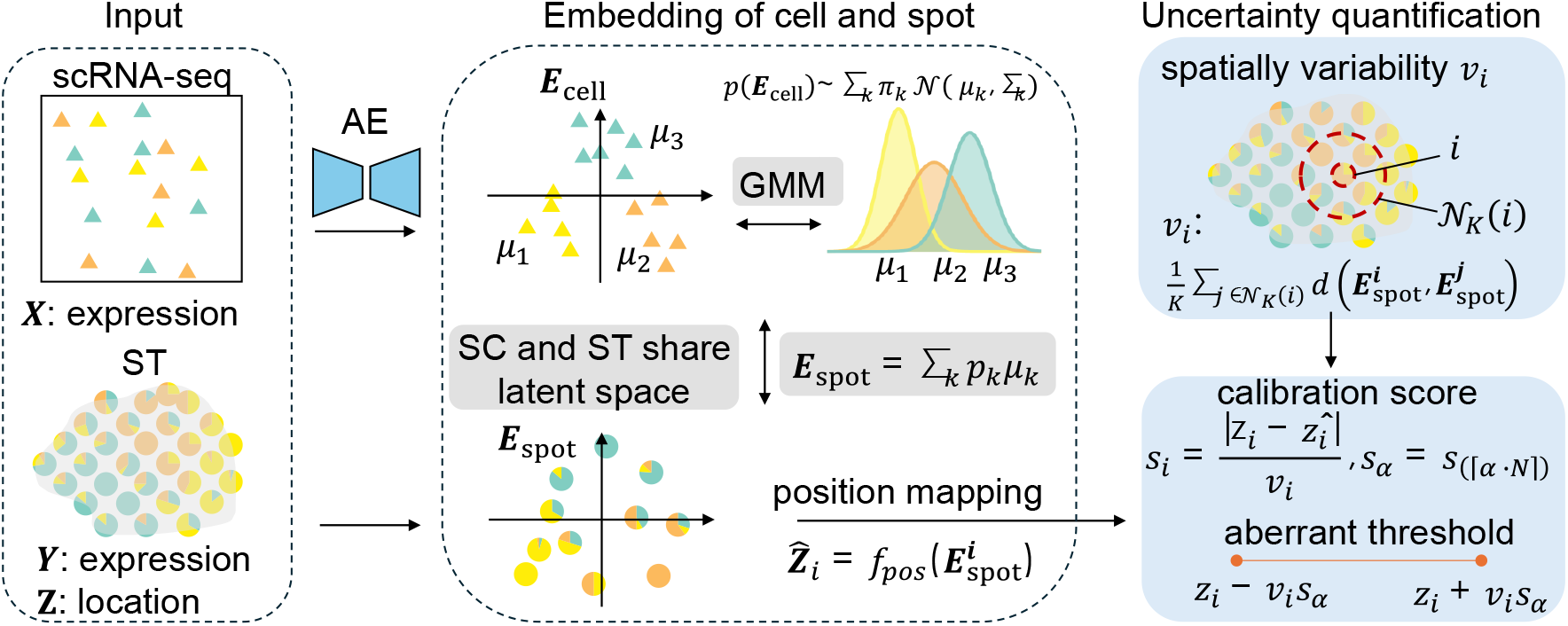
Overview of the SPADE framework. The figure is divided into three main parts: Input, Embedding, and Uncertainty Quantification. In the Input section, both scRNA-seq and ST data are provided as inputs. In the Embedding section, cells are embedded using a Gaussian Mixture Model (GMM), and each spatial spot is represented as a weighted sum of the GMM components. These spot embeddings are then used as features to predict the spatial locations of spots, yielding a positional mapping. Finally, in the Uncertainty Quantification section, a conformal prediction-based approach is applied to generate a normal interval for the predicted location of each spot. Spots whose predicted positions fall outside this interval are subsequently defined as aberrant spots.

SPADE first trains separate autoencoders on scRNA-seq and ST expression matrices to embed individual cells and spots into latent spaces of equal dimensionality, thereby achieving a unified representation across data modalities. Rather than assuming a single Gaussian distribution, SPADE models the latent distribution of scRNA-seq cells as a Gaussian mixture to capture the intrinsic complex cell-type architecture. Each cell type embedding is represented by the corresponding learned Gaussian component in the latent space. For the ST data, each spatial spot is modeled as a non-negative sparse mixture of these cell-type embeddings, reflecting the cellular composition of multi-cellular spots. This design jointly anchors cells and spots into a biologically interpretable, shared latent space. After obtaining the embeddings, we train a multilayer perceptron (MLP) that maps latent embeddings of ST spots to their observed physical coordinates, allowing the model to learn the underlying spatial organization principles encoded in healthy tissue architecture. Once trained, the mapping can predict the expected spatial position of any embedding, including cells lacking direct spatial measurements.

SPADE quantifies the uncertainty of spatial predictions using a conformal prediction scheme, yielding calibrated prediction intervals for each spot. Since there is a sense that the large difference in predicted gene expression between neighboring cells would indicate low prediction accuracy. And highly similar predicted gene expressions between neighboring cells would signify high predictive performance for the spatial location prediction method. To quantify this intuition, we introduce the spot-entered variability measure–*λ*_*i*_ by the average of embedding distances of the *k* nearest neighbors based on the spatial location. The spatial variability score is used to normalize the prediction error. Spots whose true spatial position deviates substantially from their predicted position, beyond the calibrated uncertainty bound, are classified as spatially aberrant. This uncertainty-aware calibration enables statistically principled detection of aberrant cells without reliance on manually tuned arbitrary thresholds.

### Performance evaluation using in silico data

To assess the capability of SPADE in identifying spatially aberrant spots, we generated simulated spatial transcriptomics (ST) data with a defined ground truth of aberrant spots. In detail, we perturbed the spatial coordinates of a subset of spots in a reference ST (original data) to mimic abnormal spatial displacements (in silico data).

In this simulation, 5% of spots were randomly selected and reassigned to new locations at a fixed distance from their original positions, thereby introducing spatial deviations representative of aberrant behavior. These displaced spots were labeled as spatially aberrant. Applying this procedure to the anterior Sagittal Slice of mouse brain dataset (MB_Ant) (Fig. 2A) from 10x genomics (see Methods for details) comprising 2,686 spots[22], resulted in an in silico dataset containing 127 annotated aberrant spots (Fig. 2B).We first evaluated SPADE’s spatial coordinate reconstruction on the original dataset. Normalized to [0,1], the RMSE for x and y were 0.012 and 0.019. The model achieved 90% prediction accuracy at a Euclidean distance threshold of 0.05, which increased to 95% at 0.07 (Fig. 2C), confirming its ability to accurately recover spatial organization.

We next applied SPADE to predict spatially aberrant spots using the in silico STs dataset. The nonconformity scores exhibited higher values near the tissue margins compared to the central region, indicating greater spatial uncertainty at the edges (Fig. 2D). Moreover, darker regions in the tissue H&E image corresponded to elevated nonconformity scores, suggesting that these values may capture biologically relevant or pathological variations. Consistently, prediction errors were notably higher at the predefined spatially aberrant spots (Fig. 2E). SPADE successfully identified 121 aberrant spots with an overall accuracy of 95.0% (115 out of 121), a false-positive rate of 5.96% (6 out of 121), and a recall of 90.55% (145 out of 127) (Fig. 2F). To further evaluate performance, we calculated the area under the precision–recall curve (AUPR) and the area under the receiver operating characteristic curve (AUC) across varying confidence thresholds, achieving an AUC of and an AUPR of, as shown in Fig. 2G and H. We compared SPADE against several baseline methods, including RCTD[23], CARD[24], Tangram[25], and Isolation Forest (IF)[26]. SPADE consistently achieved the best performance across all competitors, as measured by both AUROC and AUPR. To further evaluate its utility as a filtering tool for biological experiments, we assessed the ranking performance of each method using recall, precision, and F1-score across different top□K thresholds (Fig. 2I, J, and K). SPADE consistently outperformed all baseline methods across all ranking thresholds, demonstrating its robustness and practical value for prioritizing candidates in experimental validation.

To gain an understanding of parameter sensitivity, we systematically evaluated the effects of different loss weight, number of Gaussian mixture components, cell □type composition mismatches, and the model components on the model. We calculated the performance metrics, including AUROC and AUPR, between the predicted and ground truth. We varied GMM loss weight *α* to (0.05, 0.1, 1), deconvolution loss weight *β* (0.1, 0.5, 1), and Leiden resolution (0.1, 0.3, 0.5, 0.8, 1.0). We observed that AUROC remained relatively stable across most parameter configurations, while changes in Leiden resolution and *β* had a more pronounced effect on AUPR (ranged from approximately 0.52 to 0.74, Table S1). To evaluate the sensitivity of the model to cell□type mismatches between modalities, we manually removed 1 to 8 cell types (26 in total) from the scRNA□seq reference. Supplementary Table 2 summarizes the results of removing cell types. Notably, even after removing up to 8 cell types, AUROC stayed consistently high (0.901–0.915), while AUPR showed a gradual decline from 0.745 to 0.575. Finally, we performed ablation studies to assess the individual contributions of the two core components—the GMM □based VAE and the Conformal Prediction module. Removing either component led to a clear drop in performance, with AUPR decreasing from 0.745 (full model) to 0.676 (w/o CP) and 0.712 (w/o GMM □based VAE), confirming their synergistic effects (Table S3). In addition, we show the training loss curves over 1,000 epochs (Fig. S4) and report the runtime and memory usage (Table S4).

To further evaluate SPADE’s capacity to detect perturbations in gene expression programs, we designed an additional simulation experiment in which spots with altered transcriptional profiles were artificially introduced. Specifically, 5% of spots were randomly selected as ground-truth aberrant spots, and for each, we silenced 1,200, 1,800, 2,400, or 3,000 expressed genes in silico. We then applied SPADE to these perturbed datasets and assessed its ability to recover the introduced aberrations. The resulting ROC and PR curves demonstrate that SPADE consistently achieves high discrimination performance across all perturbation levels, with odds ratios ranging from 4.0 to 10.8 relative to random predictions(Fig. 2M and L).

### SPADE detected spatially aberrant spots on human SCC dataset

The tumor microenvironment (TME) provides a particularly compelling scenario to study aberrant spots [27]. Unlike normal tissues, the TME is characterized by highly heterogeneous cellular compositions, dynamic cell-cell interactions, and complex spatial organization. Cancer cells often undergo profound genetic and epigenetic alterations, leading to aberrant gene expression programs and disrupted spatial positioning within the tissue. Moreover, tumor-associated stromal cells, immune cells, and other infiltrating cells can also adopt aberrant phenotypes and spatial distributions in response to tumor-derived signals. These aberrant spots frequently contribute to cancer progression, immune evasion, and therapeutic resistance. To explore these aberrant spatial patterns, we applied SPADE to paired single-cell RNA-seq and spatial transcriptomics (ST) datasets derived from human squamous cell carcinoma (SCC) tissue (Fig. 3A). We detected 8 spatial domains by SpaGCN in this tissue (Fig. 3B). The variability in the lower-right region (domian 8) of the tissue was notably higher, suggesting an increased transcriptional heterogeneity in this area (Fig. 3C and D). After normalizing spatial coordinates to the [0, 1] range, the Euclidean distances between predicted and ground truth locations among spots spanned 0 to 0.15 (Fig. 3E).Using SPADE, we identified 22 spatially aberrant spots, predominantly localized within spatial domains 2 and 6. Domain 2 corresponded to a region with a high tumor-specific keratin (TSK) score, defined by the mean expression of the TSK gene set (MMP10, PTHLH, LAMC2, SLITRK6) [28]. Differential gene expression analysis further revealed four genes specifically enriched in aberrant spots within domain 6: TMSB15B, YIF1B, EIF2AK1, and FRYL. Among these, EIF2AK1 acts at the level of translation initiation to suppress global protein synthesis in response to cellular stress [29], while TMSB15B is implicated in cytoskeletal organization and cell migration [30]. Supplementary Figure 1 shows the expression changes of selected differentially expressed genes between the aberrant spots and their neighboring spots. To evaluate the efficiency of SPADE, we compared its spatial location mapping accuracy with two state-of-theart methods, STALocator [31] and scSpace [32]. Accuracy was quantified by varying the prediction error cutoff and computing the proportion of correctly mapped locations across thresholds(Fig. 3H). Across all cutoff values, SPADE consistently achieved the highest accuracy, demonstrating the superior reliability of its framework.

**Fig. 3.**
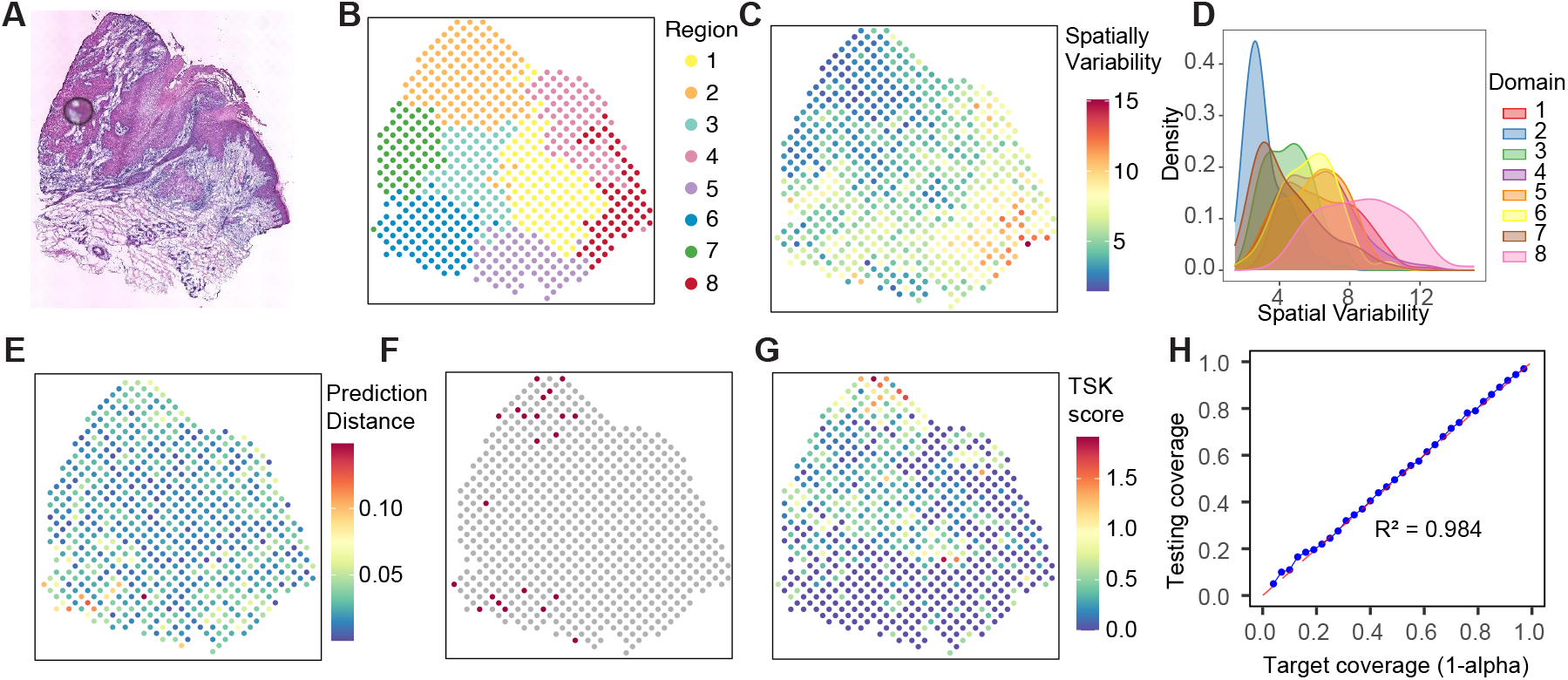
Systematic identification of spatially aberrant spots in human squamous cell carcinoma (SCC). **A.**Histological image of SCC tissue. **B**. Eight spatial domains identified by SpaGCN. **C**. Spatial variability scores computed across the SCC dataset. **D**. Distribution of spatial variability scores across the eight domains defined in (B). **E**. Prediction errors represented as the Euclidean distance between predicted and ground-truth coordinates. **F**. Spatially aberrant spots detected by SPADE. **G**. Tumor-specific keratin (TSK) score mapped across the SCC tissue section. **H**. Comparison of spot location prediction accuracy among methods across varying prediction error thresholds.

### SPADE maps single-cell transcriptomes to spatial tissue architecture

In addition to identifying spatial aberrant spots, SPADE also performs cell-type identification and spot deconvolution.In the SCC dataset, SPADE identifies transcriptionally distinct cellular populations using the Leiden clustering algorithm (21 clusters; n = 21) based on the principal component analysis (PCA) of highly variable genes [33]. The centroids of these clusters are subsequently used to initialize the Gaussian mixture model (GMM) components during model training. A key output of SPADE is the cell-type embedding, represented by the centers of the GMM components. These embeddings are well dispersed across cell types, effectively capturing the intrinsic transcriptional structure of the cellular landscape (Fig. 5A).

Each cell is assigned to the cluster with the highest posterior probability according to the GMM, and clusters containing fewer than 50 cells are removed. Following this filtering step, 13 robust clusters were retained, with clusters 5 and 8 merged based on their transcriptional similarity (Fig. 5B). All clusters were annotated to well-defined cell types using hypergeometric enrichment tests. The resulting cell-type assignments were as follows: cluster 0 (macrophages), cluster 4 (melanocytes), cluster 6 (fibroblasts), cluster 12 (myeloid-derived suppressor cells, MDSCs), cluster 14 (multiplets), cluster 16 (CLEC9A+ dendritic cells), cluster 18 (endothelial cells). The remaining clusters represented mixed or unclassified cellular populations. Compared with the original SCC histology image (Fig. 3A), the spatial boundaries of deconvolution inferred by SPADE (Fig. 5C) exhibit clear and biologically coherent correspondence with histological structure.

In addition, SPADE enables prediction of single-cell locations by projecting both cells and spots into a shared latent space. Using this framework, we mapped T cells from the scRNA-seq data onto the spatial tissue architecture. The predicted T cell distribution was highly consistent with the expression of the canonical T cell marker CD2 (Fig. 5D and E), demonstrating SPADE’s capacity to accurately reconstruct cell-type localization within the tissue microenvironment.

### SPADE identifies aberrant cell types and facilitates biological discovery

To evaluate the performance of SPADE on annotated data, we applied it to three publicly available datasets with ground-truth annotations: Drosophila embryo (DE) [34], human pluripotent stem cell (hPSC)-derived organoid [35], and human breast cancer (BCAS) [11]. For each dataset, we identified aberrant cell types by detecting the spatial aggregation of aberrant spots and examined whether the detected spots were significantly enriched within known pathological or anomalous tissue regions, as confirmed by either original annotations or literature-based evidence.

To investigate the mechanistic drivers of embryo development of the model organism Drosophila, we collected scRNA□seq and scStereo□seq data across 30 time points spanning 0 to 20.54 hours of development. These data were analyzed using SPADE in conjunction with downstream analytical pipelines. We focused on two developmental stages, 8.84 h and 17.34 h (Fig. 5A and B), where the rate of morphological change was particularly pronounced and concentrated in specific regions. At each of these two time points, we applied SPADE and identified 1,416 aberrant spots out of 28,103 spots (8.84 h) and 1,315 aberrant spots out of 25,755 spots (17.34 h) (Fig. 5C and D). At the 8.84 h stage, aberrant spots were significantly enriched in cell types corresponding to the trachea, epidermis, visceral branch, and foregut (Fig. 5E), consistent with previous reports that this period coincides with tracheal tubulogenesis, digestive system assembly, and epidermal and head remodeling [36]. At the 17.34 h stage, aberrant spots were preferentially localized to epidermal outer, fat body, epidermal dorsal, and somatic muscle cell types (Fig. 5F), aligning with the known terminal maturation of the tracheal and muscular systems, as well as the refinement of the digestive and epidermal tissues [37].

Focusing on the 8.84 h time point, we performed differential gene expression analysis between aberrant and non□aberrant spots specifically within the tracheal lineage. Genes upregulated in aberrant spots were functionally enriched in terms such as mRNA 5^′^□UTR binding, rRNA binding, cell size and cell□cycle regulation, and fatty acid *β*□oxidation (Fig. 5G to I). These enrichment patterns suggest that the aberrant cells exhibit enhanced protein transcription and translation capacity, active cell migration, and elevated energy demands, collectively underscoring their heightened metabolic and biosynthetic activity during this critical developmental window.

The BCAS tissue can be broadly classified into four major categories, as shown in Supplementary Figure 2A: tumor (DCIS/LCIS 1–5), healthy (Healthy 1–2), invasive carcinoma (IDC 1–7), and surrounding tumor (Tumor_edge 1–6). Notably, SPADE, Tangram, and Isolation Forest tended to detect spots located in both tumor and invasive carcinoma regions (Supplementary Figure 2B to E), whereas CARD and RCTD predominantly identified spots within the surrounding tumor regions (Supplementary Figure 2F and G). These observations suggest that different methods exhibit distinct preferences in terms of the types of pathological regions they prioritize, highlighting the complementary strengths of these approaches. Nevertheless, SPADE consistently captured biologically relevant regions, reinforcing its utility for aberrant spot detection in complex tissue architectures.

The hPSC-A1 tissue is classified into 9 cell types, as shown in Supplementary Figure 3A. Among these, Cholangio, Scar, and Edge are most relevant to PSC pathology, representing the bile duct epithelium, periductal fibrosis, and the inflammatory interface at the leading edge of tissue remodeling, respectively. We performed Fisher’s exact test based on the aberrant spots detected by SPADE (Supplementary Figure 3B) and found that the aberrant spots were significantly enriched in the Edge region (Supplementary Figure 3C). This enrichment result is consistent with the active role of the Edge region as an inflammatory and remodeling frontier in PSC, where bile duct injury and early fibrosis are most pronounced, thereby serving as a primary site of spatial disorganization [34, 38, 39]. To assess the robustness of this finding, we repeated the same analysis on an independent hPSC-C1 section and observed a consistent enrichment pattern (Supplementary Figure 3D–F), further supporting the reproducibility and generalizability of our observations.

### SPADE detects invasive tumor from Xenium data

With the rapid advancement of spatial transcriptomics, data resolution has reached unprecedented levels. The Xenium platform enables single-cell spatial transcriptomic profiling without the need for RNA-seq integration for cell-type identification. SPADE could be applied to Xenium data to identify spatially aberrant spots. Specifically, we isolated the spatial transcriptomic autoencoder module from our model to generate low-dimensional embeddings, then trained a mapping network that projects these embeddings to spatial coordinates. aberrant spots were subsequently identified using an uncertainty quantification framework.

We applied this approach to a Xenium dataset derived from human breast cancer samples[40], comprising 167,780 cells and 20 annotated cell types, including invasive tumor and ductal carcinoma in situ (DCIS1 and DCIS2) regions (Fig. 6A and B). SPADE identified 6,544 spatially aberrant spots (Fig. 4C). Analysis of the local neighborhood composition on the UMAP revealed that these cells were predominantly distributed across DCIS1, invasive tumor regions, and the unlabeled transition zones between them, as well as among CD4 T cells (Fig. 6D and E). Given that the tumor invasive front represents the most aggressive and genomically unstable region under high microenvironmental stress, these findings suggest that SPADE captures biologically meaningful spatial heterogeneity, potentially reflecting mechanisms underlying tumor invasiveness.

**Fig. 4.**
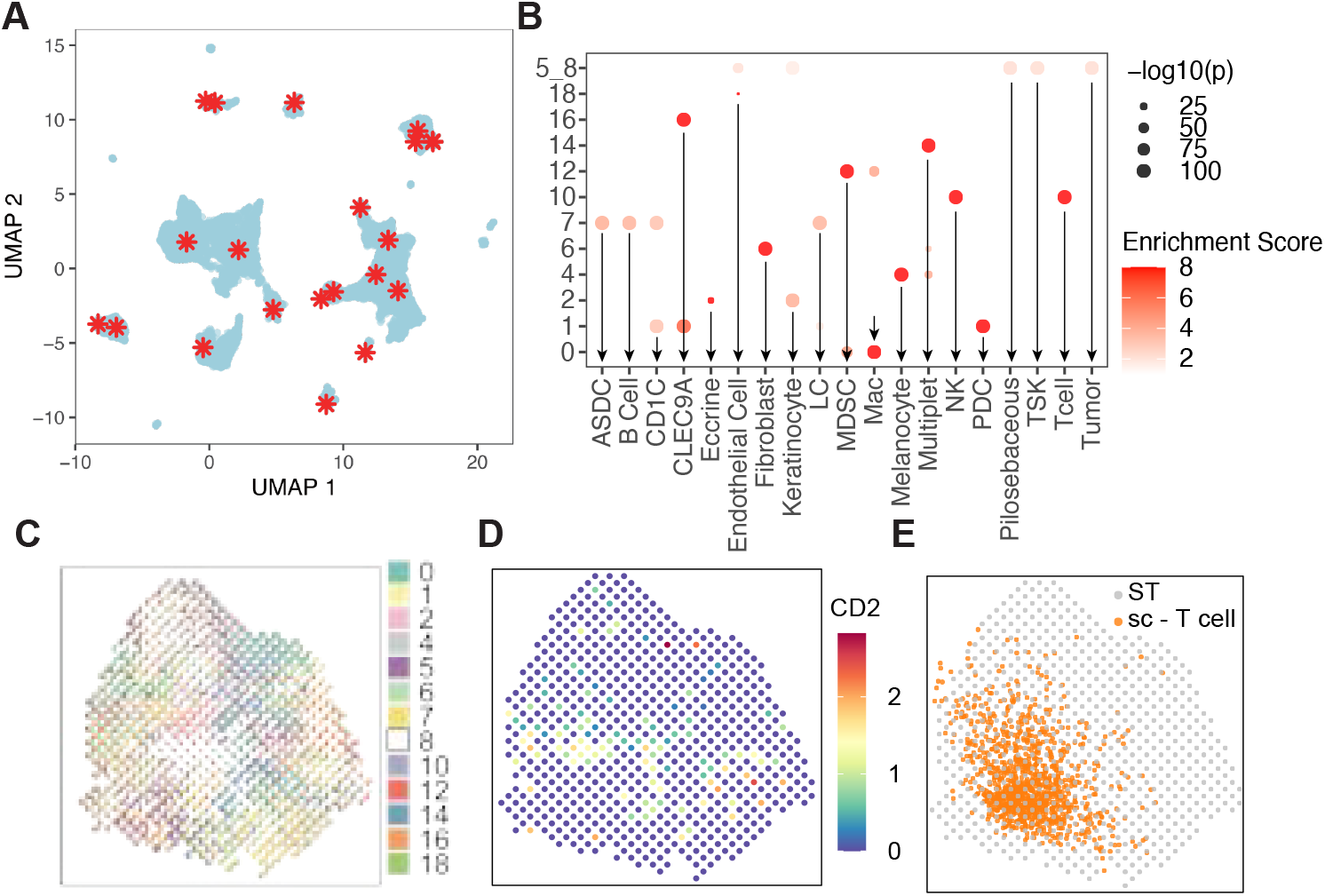
Performance of spatially aberrant cell detection in squamous cell carcinoma (SCC). A. UMAP visualization of single-cell embeddings. Red stars denote the cell-type embeddings, corresponding to the centers of GMM components. **B**. Cell-type annotations of SPADE-derived clusters. Cell-type labels were assigned based on the original study. **C**. Spot-level deconvolution results generated by SPADE. **D**. Spatial distribution of CD2 expression across spots. **E**. SPADE-predicted spatial localization of T cells from the scRNA-seq data.

**Fig. 4a.**
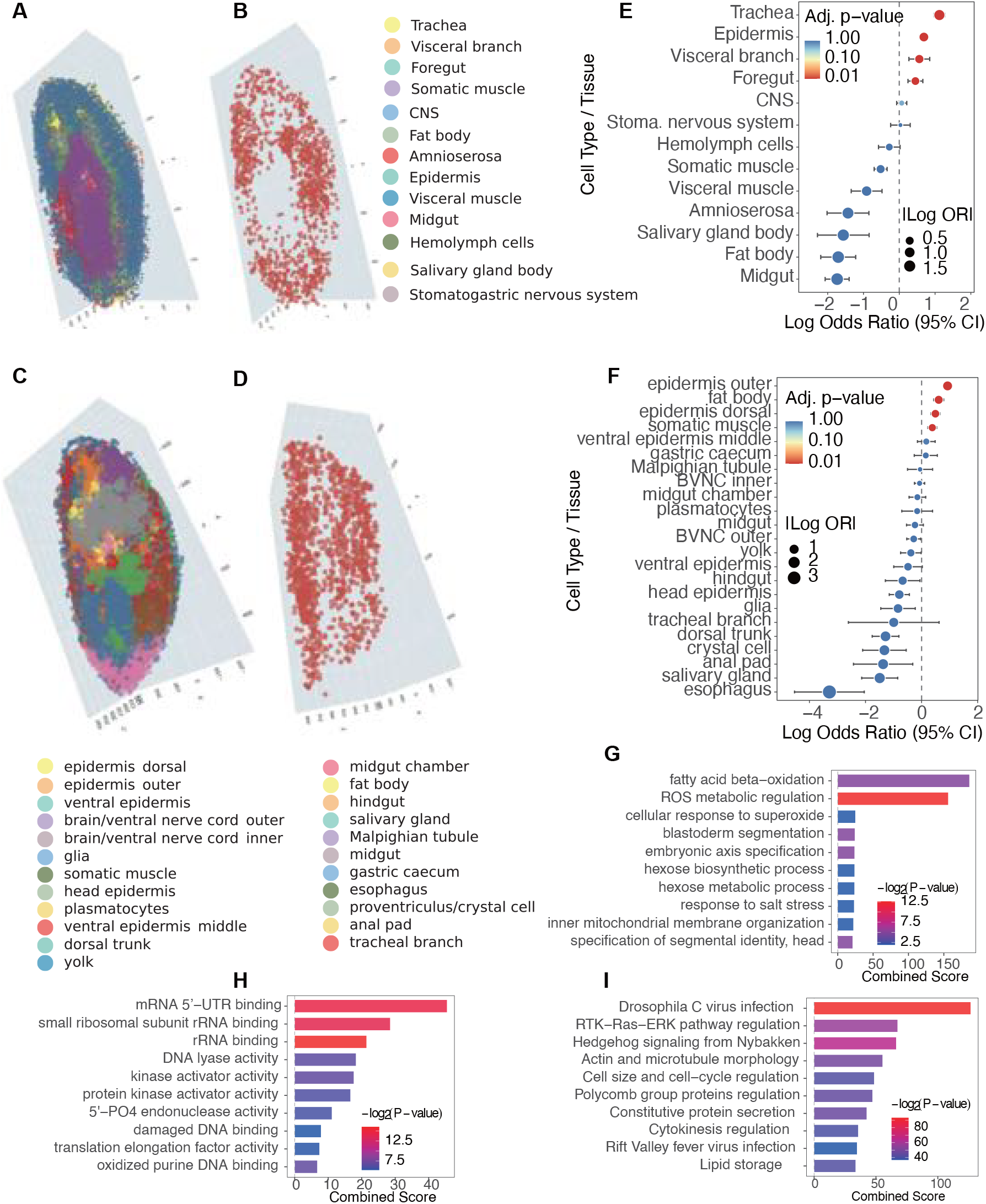
Functional and pathological characterization of aberrant spots of drosophila embryo identified by SPADE. **A**, 3D Spatial map of DE data colored by cell type of DE_8.84h. **B**, 3D Spatial map of aberrant spots detected by SPADE of DE_8.84h. **C** and **D**. 3D Spatial map of DE cell type and aberrant spots of DE_17.34h, respectively. **E** and **F**, Point plots summarizing Fisher’s exact test results for 8.84 h and 17.34 h, respectively. **G** to **I**, GO enrichment analysis of DEGs between aberrant and non-aberrant spots in the tracheal lineage.

**Fig. 5.**
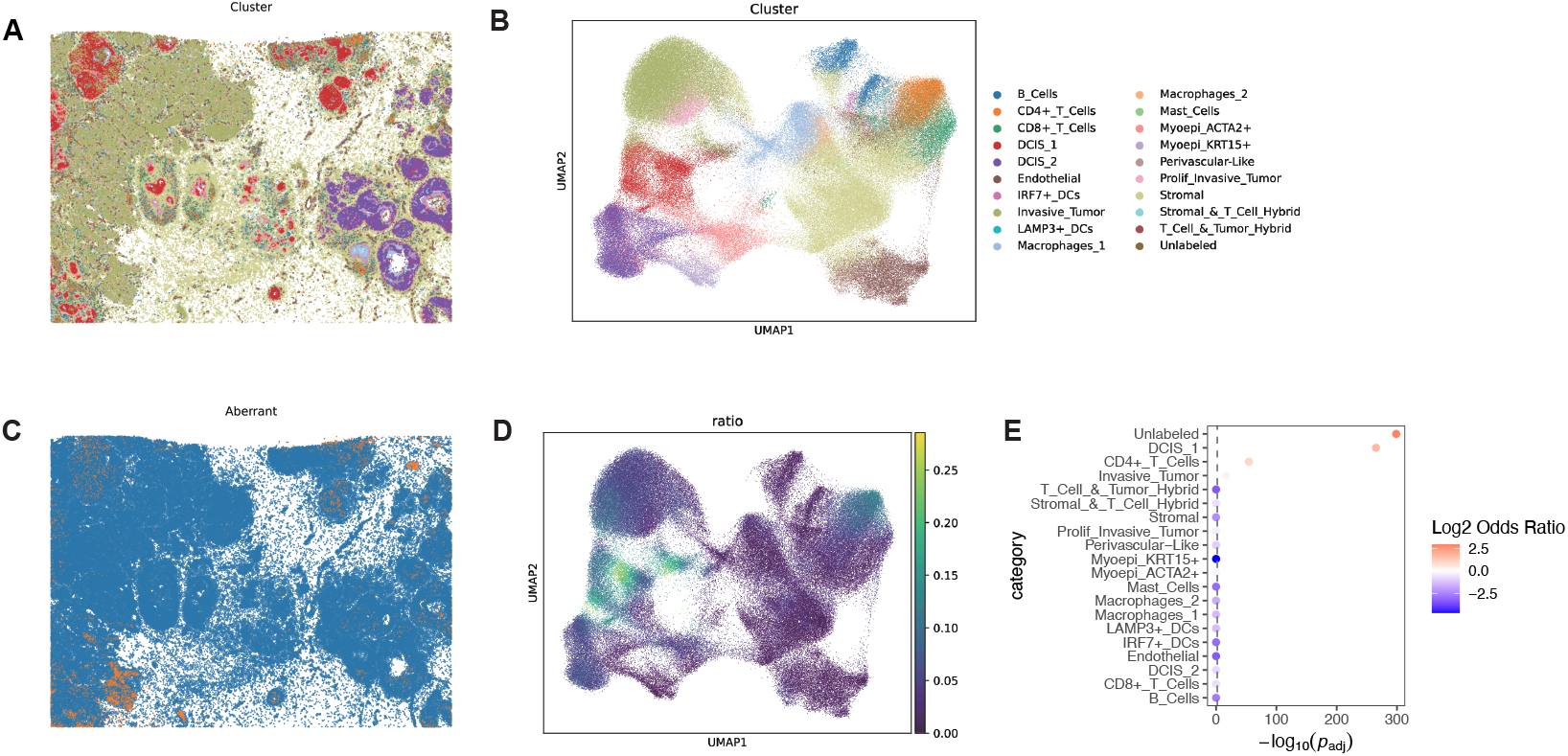
SPADE detects spatially aberrant spots in human breast cancer Xenium data. A. Histological image of the human breast cancer Xenium dataset. **B**. UMAP visualization of the Xenium data generated using Scanpy. **C**. Spatially aberrant spots identified by SPADE. Orange points indicate aberrant spots, and blue points represent other cells. **D**. Density distribution of aberrant spots in the UMAP coordinate space. **E**. Enrichment of aberrant spots across cell types. The color of each bubble represents the odds ratio of aberrant spot enrichment for each cell type relative to others, while the x-axis indicates the ‐ log_10_(*P*) value derived from Fisher’s exact test.

## Conclusion

Spatial aberrant spots, those exhibiting abnormal gene expression patterns and deviating from typical tissue organization, play critical roles in disease progression, particularly in cancer, autoimmune disorders, and neurodegenerative diseases. Although spatial transcriptomics (ST) technologies now enable the integration of gene expression with spatial context, identifying aberrant cellular states within this spatiotemporal framework remains a major computational challenge. Current methods predominantly focus on macro-scale structural modeling, with limited tools specifically designed for the detection of spatially aberrant spots.

In this work, we address the urgent need for precise, AI-driven approaches capable of high-resolution identification of such aberrant structures. By leveraging the integration of single-cell RNA sequencing (scRNA-seq) and spatial transcriptomics data, we propose a robust multimodal frame-work to quantitatively define and detect spatial aberrancy. Specifically, we map scRNA-seq cells and ST spots into a shared latent space using a variational autoencoder (VAE), and model the underlying cell type structure through a Gaussian mixture model (GMM). Each ST spot is further decomposed into a weighted combination of GMM centers via non-negative sparse regression, enabling effective deconvolution and mitigating batch effects between scRNA-seq and ST datasets. Based on the learned embeddings, we establish a mapping from spots to spatial locations. By incorporating a conformal prediction strategy, we provide well-calibrated prediction intervals for each spot’s expected spatial position, thereby enabling statistically principled identification of aberrant spots.

This study not only advances our understanding of disease mechanisms but also opens new avenues for precision medicine and the development of targeted cell and gene therapies. Future improvements will focus on enhancing the sensitivity and specificity of aberrant cell detection, as well as exploring broader biological applications, including tumor evolution, tissue regeneration, and immune infiltration, where complex spatial reorganization plays a pivotal role. The development of such spatially resolved methodologies is essential for transitioning life sciences from static molecular atlases to dynamic spatiotemporal regulatory networks.

## Method

### Data Preprocessing

To address these data quality issues, a preprocessing pipeline is proposed, consisting of two sequential steps. First, data augmentation is performed using TESLA [41], which serves dual purposes: 1) increasing the number of sample points through interpolation, and 2) imputing missing values for originally measured data, where certain true positive signals are obscured by technical noise. Subsequently, feature selection is conducted on the augmented dataset using SpaGCN [42] to identify the most spatially informative features.

### SPADE Model

We denote gene expression matrix of the scRNA-seq data as 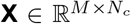 where *M* denotes the number of genes and *N*_*c*_ denotes the number of cells. 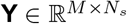 is the gene expression matrix of ST data, where *N*_*s*_ denotes the number of spots. 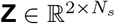 is the 2D coordinate of the location of spots.

Given the qualified gene expression matrix 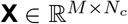 for scRNA-seq data and 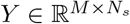 for ST data, our goal is to detect the aberrant spots with aberrant location. To represent the cell identity, we first project the spots and cells to a shared low-dimensional space by VAE-based model. To achieve cell-type deconvolution for each spot and to mitigate batch effects, we model the scRNA-seq cell embeddings with a Gaussian mixture model (GMM), such that each spot embedding can be expressed as a weighted combination of cell-type embeddings. These objectives are jointly optimized by incorporating additional loss terms into the VAE training process. Subsequently, we train a multilayer perceptron (MLP) to map each spot embedding to its corresponding spatial coordinate. To quantify positional uncertainty, we employ conformal prediction to generate prediction intervals for each inferred location. Spots whose observed coordinates fall outside the predicted intervals are defined as aberrant, reflecting abnormal spatial or transcriptional organization within the tissue.

### Cell Embedding

For the scRNA-seq data 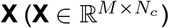 we define an encoder function *f*_cell_(·) which projects **X** into a latent space 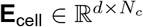 where *d < M* :

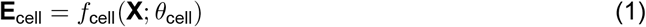

where *θ*_cell_ represents the parameters of the encoder, and **E**_cell_ is the cell embedding. This embedding is required to follow a Gaussian Mixture Distribution. We model the cell embedding **e**^cell^_*j*_ ∈ ℝ^*d*^ as a GMM with *K* Guassian distributions,

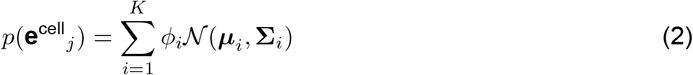

where ***µ***_*i*_ and **Σ**_*i*_ represent the mean and the Covariance matrix of the *i*-th distribution, respectively.

### Spot Embedding

Similarly, for the ST data **Y**, we use another encoder function *f*_spot_(·) to project it into a latent space 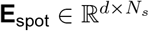

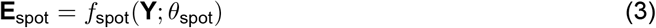

The spot embeddings **e**^spot^ ∈ ℝ^*d*^ are designed to approximate a linear combination of the Gaussian components in the GMM used for the cell embeddings.

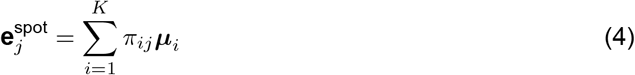

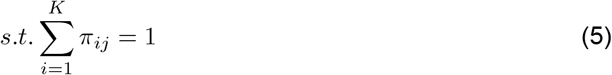

### Reconstruction with Shared Decoder

Both **E**_cell_ and **E**_spot_ are then passed through a shared decoder *g*(·) to reconstruct the original inputs **X** and **Y**:

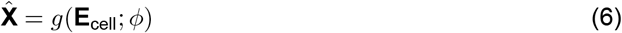

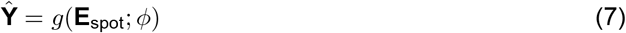

where *ϕ* denotes the parameters of the shared decoder.

### Spatial Projection

We map the spot embeddings onto their spatial coordinates **Z**:

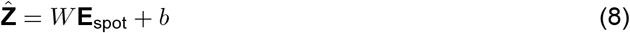

where *W* ∈ ℝ^2×*d*^ and *b* ∈ ℝ^2^ are the weights and bias terms of the linear projection.

### Conformal Prediction

After we select the tuning parameter and train the model, we can get the predicted spatial location. Furthermore, our method introduces the confidence interval using a conformal prediction approach [21, 43, 44], which can provide distribution-free uncertainty quantification. An important aspect of conformal prediction is nonconformity measure, defining how the sample differs from the rest of the data [45] under a confidence level. We describe the detailed method in the following part.

We randomly split the spots into disjoint test and calibration subsets, *S*_test_ and *S*_calib_, such that *S*_test_ ∩*S*_calib_ = ∅ and *S*_test_ ∪ *S*_calib_ = {1, 2, …, *N*_*c*_}. Then, we use test data to generate the nonconformity score, which is a normalized prediction error.

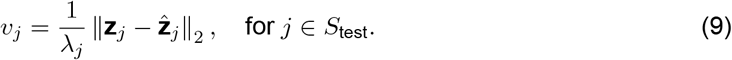

Here, we normalize the prediction error by *λ*_*i*_ to account for heterogeneous tissue structures.

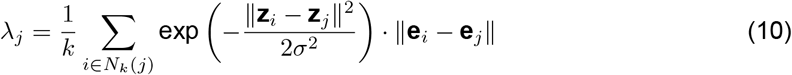

where *w_ij_* = exp 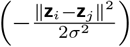 is a spatial weight and *N*_*k*_ (*j*) is the *k* nearest neighbors of spot *j*. A larger value of *λ*_*i*_ indicates that the environment around the point is more variable or that the point is in a less common location, which may imply lower prediction accuracy.

Then we get 95% quantile of the nonconformity score:

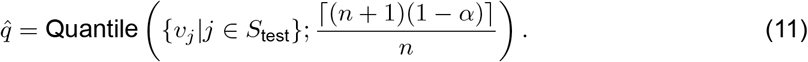

Where n is the number of samples. *α* is the quantile threshold which is usually chosen as 0.1 or 0.05. Then in calibration data samples, we get 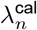 similar with getting them on testing samples and then get the prediction interval/size:

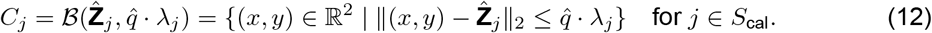

The smaller the interval/size, the more accurate results the method gets. For spot *j*, we identify it as aberrant if it is not in the confident interval:

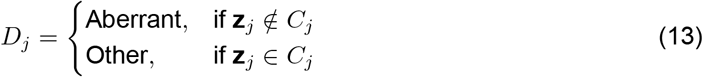

To get full prediction interval/size including size in testing data, we switch testing data and calibration data. Specifically, we get *v*_*j*_ in equation 9 from calibration data and get final size in equation 12 from testing data.

### Loss function

The overall loss function is:

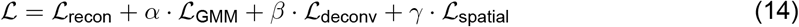

where the individual components of the loss function are defined as follows:

- **Reconstruction Loss(ℒ** _**recon**_): This term measures the reconstruction error between the original gene expression matrix and its reconstructed outputs:

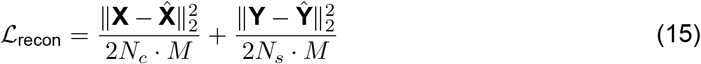
- **GMM Fitting Loss (ℒ** _**GMM**_): This term ensures that the cell embeddings follow a Gaussian Mixture Model (GMM) by minimizing the negative log-likelihood:

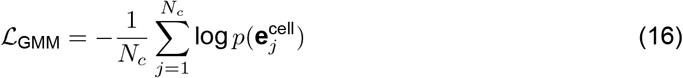

where 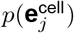 is given by:

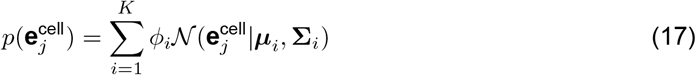
- **Deconvolution Loss (ℒ** _**deconv**_): This term ensures the spot embedding is a combination of the cell-type embedding.

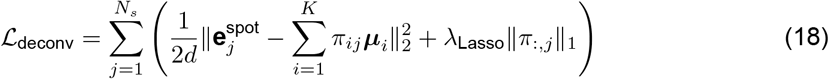 Since a single spot in spatial transcriptomics typically contains cells of the same type or a few closely related cell types, which correspond to a sparse *P* matrix. Here, we add Lasso (*l*_1_ regularization) to the loss function for sparsity.
- **Spatial Projection Loss (ℒ**_**spatial**_): This term measures the discrepancy between the predicted spatial coordinates 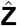 and the true spatial coordinates **Z**. Here, we use a Huber loss, which is inherently more robust to outliers and reduces the influence of noisy or structurally aberrant spots during training. The loss of *j*-th spot is defined by:

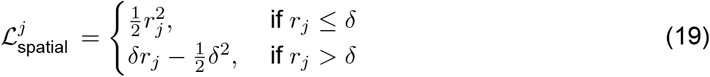

Where 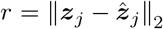 is the Euclidean norm of the residual vector between the learned embedding ***z***_*j*_ and its predicted spatial location 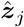 and *σ* set to the 95th percentile of the empirical distribution of all *r*_*j*_ values computed over the current training batch.

The hyperparameters *α, β*, and *γ* control the relative importance of each loss component.

## Data and code availability

The mouse brain tissue ST data used during this study was downloaded from the 10× Genomics website (https://www.10xgenomics.com/datasets/mouse-brain-serial-section-2-sagittal-anterior-1-standard). Mouse brain scRNA-seq data was downloaded from the 10× Genomics website (https://www.10xgenomics.com/datasets/10k-mouse-forebrain-ffpe-tissue-dissociated-using-gentlemacs-dissociator-singleplex-sample-1-standard). Human SCC datasets, Gene Expression Omnibus (GEO): GSE144240. The source code can be found here: https://github.com/sohu-123-lang/SPADE/blob/main/README.md.

## Ethical statement

This study involved secondary analysis of publicly available and de-identified datasets. No new human participants were recruited, no new human samples were collected, and no animal experiments were conducted by the authors. Therefore, institutional ethics approval and informed consent were not required for the work reported in this study. The ethical and data-use requirements associated with each source dataset were followed.

## Data availability

No new raw sequencing data were generated in this study. The publicly available datasets analysed in this work are described in the Methods section and are available from their original repositories. In particular, the human squamous cell carcinoma dataset is available from the Gene Expression Omnibus under accession number GSE144240. The processed files, analysis inputs, and other materials newly generated for this study have been deposited in Zenodo at https://doi.org/10.5281/zenodo.21810260 [AUTHOR CONFIRMATION REQUIRED: retain this sentence only if the Zenodo record contains all newly generated materials and is publicly accessible or has a valid reviewer-access link].

## Code availability

The source code for SPADE, documentation, and implementations used for the benchmark comparisons are available at https://github.com/zhangzheng0131/SCAD/tree/main. The code will also be deposited in NGDC BioCode at [AUTHOR CONFIRMATION REQUIRED: insert the final BioCode accession number and URL] in accordance with the journal’s data and code policy. The repository includes the software environment and instructions required to reproduce the analyses.

## CRediT author statement

**Zheng Zhang**: Conceptualization, Methodology, Software, Formal analysis, Investigation, Visualization, Writing – original draft. **Xubin Zheng**: Conceptualization, Methodology, Supervision, Writing – review & editing. **Qiuyue Yuan**: Methodology, Validation, Formal analysis, Writing – review & editing. **Rui Luo**: Conceptualization, Supervision, Project administration, Funding acquisition, Writing – review & editing.

## Acknowledgments

The authors thank the developers and contributors of the publicly available datasets and software resources used in this study. This work was supported by the Guangdong Basic and Applied Basic Research Foundation, China (No. 2026A1515060002); the National Natural Science Foundation of China (Nos. 32300554 and 62506315); the Guangdong Provincial Key Laboratory of Mathematical and Neural Dynamical Systems, China (No. 2024B1212010004); the Hong Kong Research Grants Council (CityU 21213626); the Guangdong Provincial Natural Science Foundation, China (No. 2026A1515011937); and City University of Hong Kong (No. 9610639).

## Declaration of AI and AI-assisted technologies in the writing process

During the preparation of this work, the authors used [AUTHOR CONFIRMATION REQUIRED: insert the name of the AI tool/service, if used] in order to [AUTHOR CONFIRMATION REQUIRED: state the specific purpose, e.g., improve language clarity, translate draft text, generate code, or process data]. After using this tool/service, the authors reviewed and edited the content as needed and take full responsibility for the content of the publication.

